# Transcriptomic insights into the establishment of coral-algal symbioses from the symbiont perspective

**DOI:** 10.1101/652131

**Authors:** Amin R Mohamed, Natalia Andrade, Aurelie Moya, Cheong Xin Chan, Andrew P. Negri, David G. Bourne, Eldon E Ball, David J Miller

**Affiliations:** CSIRO Agriculture and Food, Queensland Bioscience Precinct, St Lucia 4067, Queensland Australia; Zoology Department, Faculty of Science, Benha University, Benha 13518, Egypt; ARC Centre of Excellence for Coral Reef Studies, James Cook University, Townsville 4811, Queensland, Australia; Department of Molecular and Cell Biology, James Cook University, Townsville 4811, Queensland, Australia; AIMS@JCU, Australian Institute of Marine Science, Department of Molecular and Cell Biology, James Cook University, Townsville 4811, Queensland, Australia; Institute for Molecular Bioscience, The University of Queensland, Brisbane 4072 Queensland, Australia; Australian Institute of Marine Science, Townsville Queensland, Australia; Department of Marine Ecosystems and Impacts, James Cook University Townsville 4811 Queensland, Australia; Division of Ecology and Evolution, Research School of Biology, Australian National University, Acton, ACT 2601, Australia

**Author notes:** Correspondence: D Miller and A Mohamed.

**Keywords:** coral, Symbiodiniaceae, transcriptome repression, symbiosis, dual RNA-Seq

## Abstract

Despite the ecological significance of the mutualistic relationship between Symbiodiniaceae and reef-building corals, the molecular machinery underpinning the establishment of this relationship is not well understood. This is especially true of the symbiont side, as previous attempts to understand the interaction between coral larvae and Symbiodiniaceae have focused nearly exclusively on the host. In the current study, *Acropora tenuis* planula larvae were exposed to a compatible strain of Symbiodiniaceae (*Cladocopium*) and the transcriptomic landscape of the symbiont profiled at 3, 12, 48 and 72 h post-exposure using RNA-Seq. The transcriptomic response of *Cladocopium* to the symbiotic state was complex, the most obvious feature being an extensive and generalised downregulation of gene expression. Included in this “symbiosis-derived transcriptional repression” were a range of stress response and immune-related genes. In contrast, genes implicated in metabolism were upregulated in the symbiotic state. Consistent with previous ecological studies, this transcriptomic response of *Cladocopium* implied that active translocation of metabolites to the host occurred, and thus that the mutualistic relationship can be established at the larval stage. This study provides novel insights into the transcriptomic remodelling that occurs in Symbiodiniaceae, with important implications for understanding the establishment of symbiosis between corals and their dinoflagellate partners.

## Introduction

The metabolite exchange between members of the Symbiodiniaceae and their adult coral hosts has been extensively studied. In this mutualism, the coral host provides the symbiont with inorganic nutrients (Trench, 1979; Allemand et al., 1998; Leggat et al., 2003; Weis et al., 2008), whereas the symbiont can satisfy most of the energy requirements of the host in the form of organic compounds (glucose, glycerol, fatty acids and amino acids) (Muscatine 1990, Grant et al., 1997, Papina et al., 2003, Burriesci et al., 2012). Although cnidarian-Symbiodiniaceae mutualisms have been extensively investigated, little is known about this relationship during the host larval stage, particularly in relation to metabolite exchange and symbiosis-related differential gene expression. Indeed, it is unclear whether the mutualistic nature of this relationship is already set up during early life history or it becomes established at a later stage.

Genome and transcriptome data are now available for multiple Symbiodiniaceae genera (LaJeunesse et al., 2018); for example, genome assemblies have been published for *Symbiodinium* (as Clade A; Aranda et al., 2016), *Breviolum* (as *Symbiodinium* Clade B1; Shoguchi et al., 2013), *Fugacium* (as *Symbiodinium* Clade F; Lin et al. 2015, Liu et al. 2018), *Symbiodinium sensu stricto* (as Clade A: *Symbiodinium microadriaticum*; Aranda et al., 2016, Shoguchi et al., 2018), and *Cladocopium* (as *Symbiodinium* Clade C; Liu et al., 2018, Shoguchi et al., 2018) in recent years. However, to date the exploitation of these genomic resources has been primarily comparative rather than functional, hence our limited understanding of the molecular events underlying the establishment and breakdown (dysbiosis) of the symbiosis. Indeed, the few published studies on the molecular bases of coral - Symbiodiniaceae interactions have focused exclusively on the host. For example, microarray-based studies implied that infection with compatible strains of symbiont had little or no impact on transcription in coral larvae (Voolstra et al., 2009, Schnitzler and Weis 2010). A more recent RNA-Seq based study (Mohamed et al., 2016) identified a transient period of differential expression that involved approximately one thousand genes 4 h after the exposure of *Acropora digitifera* planula larvae to a competent symbiont. Symbiosis-specific gene expression has also recently been reported in the sea anemone *Exaiptasia* (Bucher et al., 2016, Wolfowicz et al., 2016) which also hosts Symbiodiniaceae. On the other hand, very few studies (reviewed in Mies et al. 2017a) have addressed symbiont gene expression, with the exception of the recent paper by Bellantuono et al. (2019) on the *Durusdinium trenchii* in *Exaiptasia*.

To date, the only candidate symbiosis-related gene on the algal side of coral-Symbiodiniaceae interactions is a H^+^-ATPase in (formerly clade A) *Symbiodinium* (Bertucci et al., 2010). Mies et al. (2017b) verified the expression of this algal H^+^-ATPase in Symbiodiniaceae infected larvae of the coral *Mussismilia hispida*. Working with cultured Symbiodiniaceae, Rosic et al. (2014) suggested that protein kinases might be involved in the establishment of symbiosis, as well as identifying the H^+^-ATPase.

To investigate symbiont responses during the transition from free-living to symbiotic states, transcriptome-wide gene expression levels in *Cladocopium* (previously known as *Symbiodinium* Clade C) were compared *in hospite* to the cultured control (representing the free-living state) at 3, 12, 48 and 72 h post-infection by mapping Illumina RNA-Seq reads onto the *Cladocopium* transcriptome. By contrast with previous studies, in this time-course infection study thousands of genes were differentially regulated in the symbiotic state compared to the free-living condition. The *Cladocopium* response to the symbiotic lifestyle was complex, one aspect of this being suppression of transcription of a wide range of genes, including those implicated in immunity and stress responses. This molecular response is consistent with the idea that the *Cladocopium* niche *in hospite* represents a very stable environment compared to the free-living condition. Against the background of genome-wide transcriptional repression, the predicted functions of the genes upregulated in the symbiotic state imply that active symbiont to host metabolite translocation is initiated at the host larval stage. This paper is the first to comprehensively document the molecular events underpinning the transition of Symbiodiniaceae algae from free-living to symbiotic states during the infection of a compatible coral host.

## Material and Methods

### Algal culture

*Cladocopium* (AIMS-aten-C1-MI-cfu-B2) was obtained from the Symbiont Culture Facility at the Australian Institute of Marine Science (AIMS) and was used in this infection experiment. This strain was originally isolated from *Acropora tenuis* collected from Magnetic Island. Cultures were grown in filtered seawater (FSW) supplemented with Daigo IMK (Wako Pure Chemical Industries, Ltd.) and maintained under a 12/12-h light/dark cycle before they were used for infecting coral larvae.

### Coral larvae and infection experiment

*Acropora tenuis* colonies were collected from Magnetic Island in November 2014 and maintained in the National Sea Simulator (SeaSim) at the Australian Institute of Marine Science (AIMS) in Townsville until spawning occurred (10 November 2014). Following fertilization embryos were raised in 0.2 µm FSW under ambient conditions. Six days post-fertilization, ∼ 1000 planulae were distributed into each of twelve 1 L plastic containers containing 1 L of 0.2 μm FSW, giving triplicates of infected coral larvae at four time points. Cultured algae were washed three times in 0.2 μm FSW and added at a density of 10^5^ cells/ml. Containers were held at ambient temperature (approximately 26°C) and light conditions (80-100 μmol photon m^-2^ s^-1^ measured using a LI-193SA Underwater Spherical Quantum Sensor with LI-250A Light Meter, LI-COR® Inc., Lincoln, NE, USA). At 3, 12, 48 and 72 h post-infection, ∼200 larvae from each replicate were collected as described previously (Mohamed et al., 2016, 2018). Briefly larvae were washed in 0.2 μm FSW (ensuring as little liquid carry-over as possible), snap-frozen and stored at −80°C until further processing. At each time point, 10 larvae were washed and inspected under a fluorescence microscope to quantify the infection as numbers of symbionts per larva.

### RNA isolation and high-throughput sequencing of messenger RNA

Total RNA was isolated from the frozen larval samples using the RNAqueous® Total RNA Isolation Kit (Ambion). Larvae were lysed twice for 20s at 4.0 m s^−1^ in Lysing Matrix D tubes (MP Biomedicals, Australia) containing 960 µL of lysis/binding solution plus 80 µL of the Plant RNA Isolation Aid (Ambion, USA) on a FastPrep®-24 Instrument (MP Biomedicals, Australia). RNA was bound to the columns, washed multiple times, eluted in 40 μL of RNase-free water and stored at −80°C. RNA was quantitated and its integrity assessed using a NanoDrop ND-1000 spectrometer (Wilmington, DE, USA) and Agilent 2100 Bioanalyzer (Santa Clara, CA, USA). Messenger RNA (mRNA) was isolated from 1 μg of total RNA and 12 RNA-Seq libraries prepared using the TruSeq RNA Sample Preparation Kit (Illumina). Libraries were sequenced on an Illumina HiSeq 2000 platform. Sequencing produced a total of ∼ 400 million individual 100 bp paired-end reads.

### RNA-Seq data analysis

When isolated, the RNA population is a mixture of both host and symbiont RNA. To separate symbiont reads from those of the host, raw Illumina reads were mapped simultaneously onto the *A. tenuis* (https://matzlab.weebly.com/data--code.html) and *Cladocopium* transcriptomes (Levin et al., 2016) using BOWTIE2 (Langmead & Salzberg 2012) with default parameters. In this paper we focus only on transcriptome changes in the algal symbiont, hence transcript abundances were estimated from libraries mapped to *Cladocopium* only using RSEM (Li & Dewey 2011). These data were compared with RNA-Seq data for cultured *Cladocopium* (Levin et al., 2016) (GEO database GSE72763) and analysed as described earlier. Importantly, the same strain of *Cladocopium* from the same source (AIMS Symbiont Culture collection) was employed as was used by Levin et al. (2016).

Differential gene expression was inferred based on the raw counts using the edgeR package (Robinson et al. 2010) in R (https://www.r-project.org/). The twelve symbiotic *Cladocopium* samples at 3, 12, 48 and 72 h post-infection were compared to the control samples. *P*-values for differential gene expression were corrected for multiple testing using the Benjamini and Hochberg algorithm. Genes that have FDR at most 0.01 and at least absolute log_2_(fold change) ≥ 1 were considered significant and used for downstream analyses. To compare expression profiles across the four infection time points, significant differentially expressed genes (DEGs) were hierarchically clustered using the TMM-normalized expression values (FPKM) that were log_2_-transformed and median-centered by transcript (Robinson & Oshlack 2010; Haas et al., 2013). To infer the functions of up- and downregulated genes, BLASTX analysis (e ≤ 10^-5^) was performed against the Swiss-Prot database. The functional profile for these annotated DEGs was inferred in a two-stage analysis as previously described (Mohamed et al. 2016, 2018). The RNA-Seq reads used in this study have been submitted to the NCBI Gene Expression Omnibus (GEO) database under Accession number GSExxxxx.

## Results

### Symbiont transcriptome landscape during host infection

Transcriptomic profiles were compared between *Cladocopium* at 3, 12, 48 and 72 h post-infection and the corresponding cultured stage as the control. Sequencing yielded an average of 33 million paired-end (PE) reads per library (Supplementary Table S1). The proportion of the symbiont reads relative to host reads increased from 0.6 to a maximum of 6.5 at the end of the infection (Supplementary Fig. S1), reflecting increasing transcriptional output during symbiont colonization (Fig.1 and Supplementary Table S1). These proportions are in the same range as in previous dual RNA-Seq of host-parasite interactions (see, for example, Choi et al., 2014, Nuss et al., 2017, LaMonte et al., 2019). Hierarchical clustering of the expression data revealed agreement among the biological replicates. Pairwise Spearman correlations among samples showed lower variability within replicates of the same treatment as compared with variation between treatments (Supplementary Fig. S3). Clustering revealed two main groupings, with all symbiotic samples in one cluster and control samples in the other. These exploratory analyses qualified the RNA-Seq data as robust and allowed downstream analyses to proceed with confidence.

**Fig. 1.**
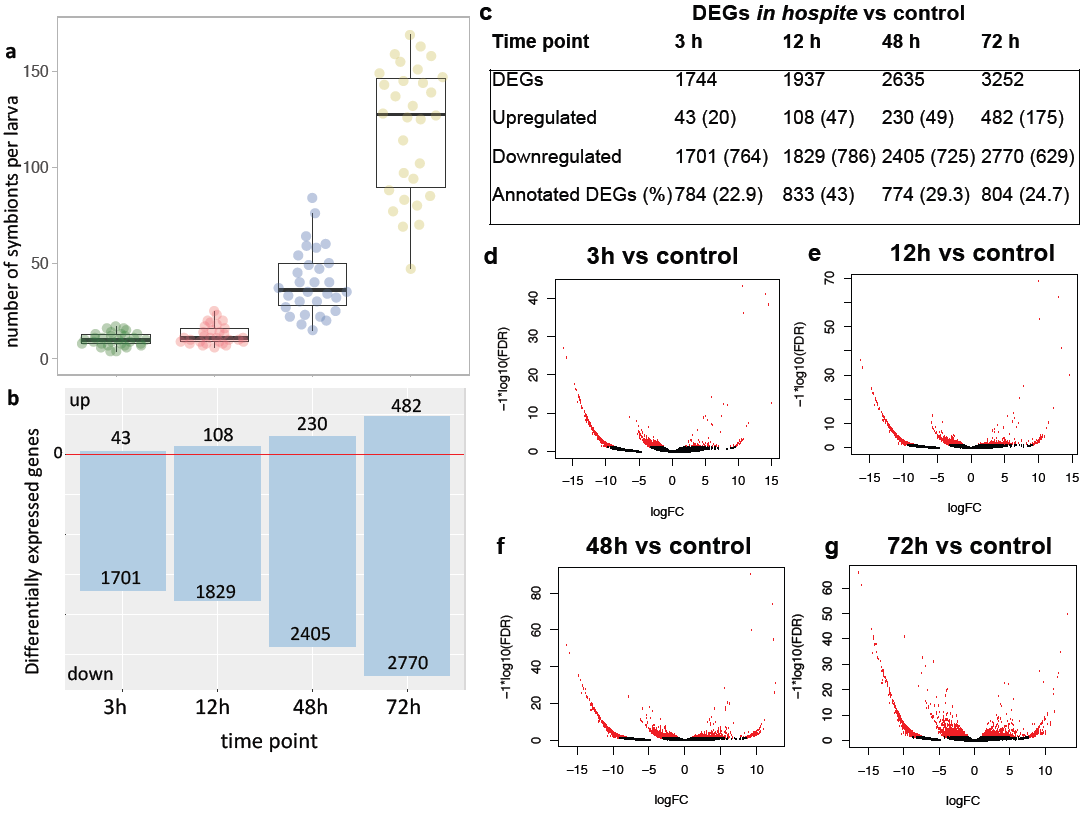
Summary of differential gene expression profiles in *Cladocopium* during uptake by *A. tenuis* larvae. **a** Boxplot showing numbers of *Cladocopium* cells within infected larvae (N=30, 3 replicates x 10 larvae/timepoint) at the times indicated. **b** Bar graph showing the numbers of differentially expressed genes in symbiotic *Cladocopium* infecting *A. tenuis* larvae compared to controls at 3, 12, 48, and 72 h. **c** Summary table for the differential expression and annotation results. An adjusted P ≤ 0.01 and E-value cut off ≤10^−5^ were used to filter differentially expressed genes and for BLASTX searches against the Swiss-Prot database. **d, e, f and g** Volcano plots showing *Cladocopium* genes differentially expressed at 3, 12, 48, 72 h during infection compared to control. The plots show false discovery rate (-log_10_FDR) as a function of log_2_(fold change). The red dots represent the significant differentially expressed transcripts.

### Differential gene expression analysis

Differential gene expression (≥ 2-fold differential expression and FDR ≤ 0.01) in symbiotic *Cladocopium* relative to controls revealed extensive suppression of gene expression throughout the infection process (Fig.1; Supplementary Table S2); although numbers of genes upregulated and downregulated increased during the infection process, the numbers downregulated far exceeded those upregulated. The range of log_2_fold changes and FDR are summarised in (Fig.1). Hierarchical clustering of the significant DEGs revealed distinctive expression profiles between symbiotic *Cladocopium* and control samples (Fig. 2). Using a corrected *P*-value ≤ 0.05, no significant GO enrichment at any of the three categories (biological process, molecular function, cellular component) could be detected among upregulated genes. For downregulated genes, significant GO enrichment was detected at each time point (Fig. 3; Supplementary Table S3) throughout the infection process, and clearly implies that protein synthesis and processing are also downregulated in *Cladocopium*, presumably as a corollary to the observed transcriptional downregulation during interaction with coral larvae.

**Fig. 2.**
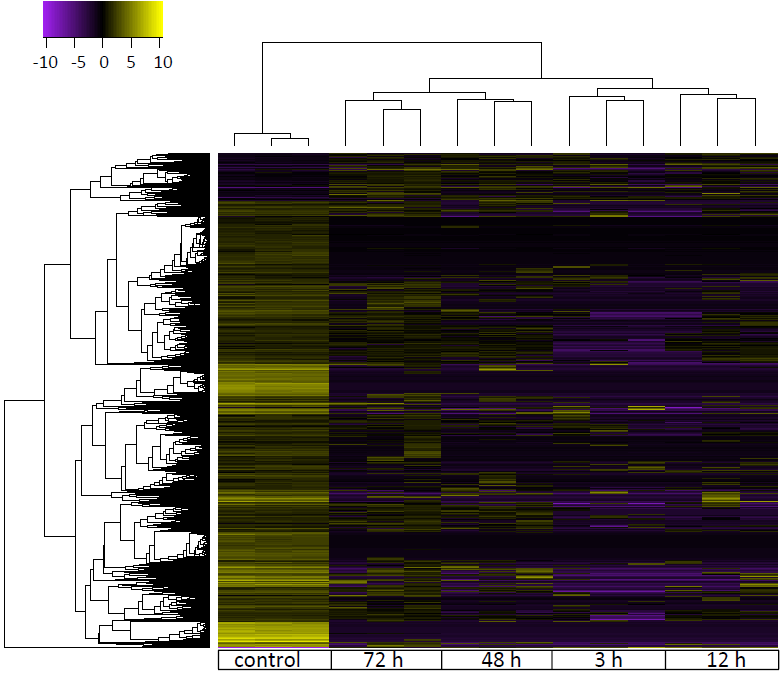
Symbiosis-derived transcriptome repression in symbiotic *Cladocopium*. Heatmap of *Cladocopium* genes with significant differential expression (fold ≥ 2, FDR ≤ 0.01) between symbiotic samples and controls at 3, 12, 48 and 72h post-infection. The hierarchical clustering obtained by comparing the expression values (Fragments Per Kilobase of transcript per Million; FPKM) for symbiotic *Cladocopium* samples compared against the control. Expression values are log2-transformed and then median-centered by transcript

**Fig. 3.**
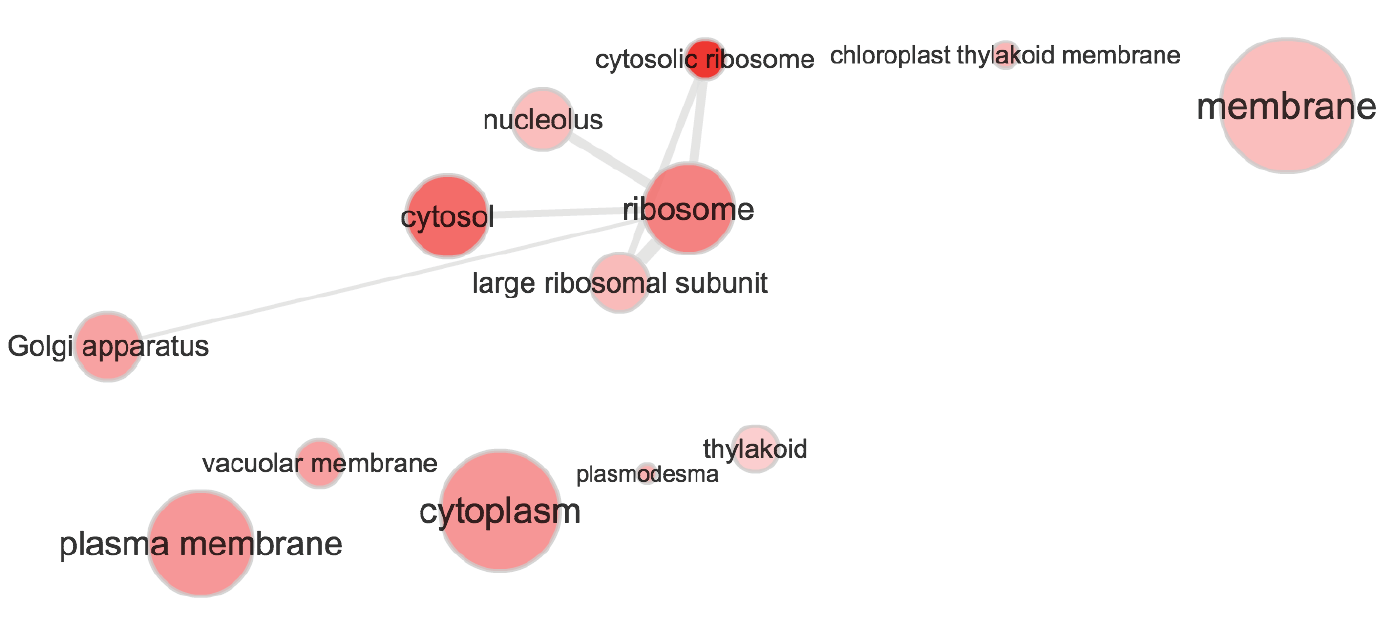
REVIGO GO relationship graph of cellular component GO terms of genes downregulated in symbiotic *Cladocopium*, using GO terms enriched with a corrected P-value ≤ 0.05. Full GO enrichment analyses results are provided in the supplementary table S3.

### Response profiles of genes with symbiosis-related functions

GO enrichment analysis provided a general overview of the molecular events occurring in *Cladocopium* during its infection of coral larvae. The GO database is biased towards processes in model organisms; consequently, another phase of data analysis was undertaken based on literature searches highlighting functions implicated in symbiosis. Significant responses were detected in categories of genes associated with stages of symbiosis establishment between *Cladocopium* and coral larvae (metabolism, transport and stress response). The composition of each of these categories of genes and others is explored in more detail below.

#### Genes implicated in the transition from free-living to symbiotic states

Analyses of *Cladocopium* responses reveal genes implicated in the transition from free-living to symbiotic lifestyle including genes coding for N-glycan biosynthesis, ankyrin (ANK) proteins, ankyrin repeat (AR) containing proteins, tetratricopeptide repeat (TPR) containing proteins, flagellar motility and motor proteins. The rationale for examining these genes in the context of the free living/symbiotic transition is as follows: N-glycans are implicated in the symbiont recognition process (Lin et al. 2000, Wood-Charlson et al. 2006), proteins containing repetitive motifs have frequently been associated with microbial infection of eukaryotic cells (see below) and flagellar motility is not required by Symbiodiniaceae *in hospite*.

Seven genes implicated in N-glycan biosynthesis were differentially expressed during the infection process (Fig. 4a, 6 and Supplementary Table S4). Most of these code for N-glycan early processing proteins such as alpha-mannosidase 1(MNS1) and alpha-1,3-mannosyl-glycoprotein beta-1,2-N acetylglucosaminyltransferase (CGL1) and were downregulated during early infection relative to controls, the exception being a mannosidase alpha class 1B member 1 (MAN1B1) that is required for N-glycan trimming being upregulated.

**Fig. 4.**
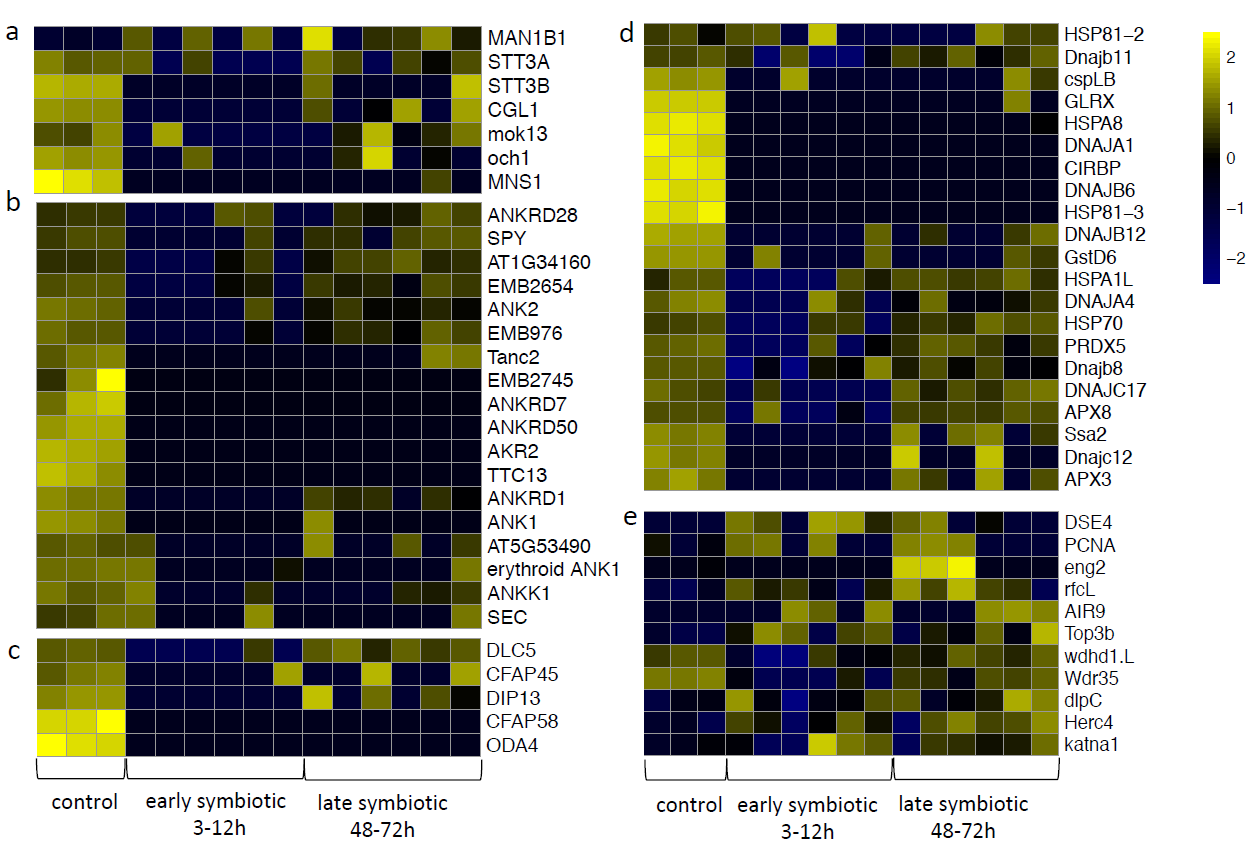
Heat maps of **a** genes implicated in N-glycan biosynthesis, **b** genes encoding ankyrin proteins/TPR domains, **c** genes implicated in flagellar motility, **d** genes implicated in stress responses, **e** genes implicated in replication in symbiotic *Cladocopium* at 3h, 12h, 48h, 72h post-infection and in controls. The hierarchical clustering shown here was obtained by comparing the expression values (fragments per kilobase of transcript per million; FPKM) for symbiotic samples and the controls. Expression values were log_2_-transformed and median centered by transcript. Relative expression levels are shown in yellow (upregulated) and purple (downregulated).

A total of 18 genes encoding ANK/AR or TPR proteins were differentially expressed during the infection process (Fig. 4b, 6 and Supplementary Table S5). With few exceptions, these genes were downregulated after infection compared to the controls, however, while some of these were upregulated to varying extents later in the infection process (eg. ANKRD28), for several others (eg. ANKRD7), no further significant expression was observed even at 72 h after infection (Fig 4d).

Perhaps not surprisingly, five genes involved in flagellar motility (Fig. 4c, 6 and Supplementary Table S6) were differentially expressed in *Cladocopium* during the establishment of symbiosis. Among these, two genes that were highly expressed in the controls, those coding for cilia and flagella associated protein 58 (CFAP58) and flagellar outer dynein arm heavy chain beta (ODA4; which is a force generating protein of eukaryotic cilia and flagella) were downregulated at all time points examined.

#### Changes in expression of genes implicated in immunity, stress responses and replication

Nine genes with immune-related functions were significantly downregulated (Fig. 6 and Supplementary Table S7) during infection including programmed cell death 2-like (PDCD2L), 3 genes encoding homologs of a key regulator of innate immune response NOD-Like Receptor C3 (NLRC3) and sushi domain containing 2 (Susd2). A total of 21 likely stress response genes were differentially expressed over the course of the infection process (Fig. 4d, 6 and Supplementary Tables S8). Five genes in this category, all of which were strongly expressed in free-living controls, were downregulated throughout the infection period examined, including an Hsp70 family member (HSPA8) and its Hsp40 co-chaperone (DNAJA1). As in the case of other functional categories, other stress response genes were downregulated early in the infection but increased their expression later in the infection process, including a distinct HSP70 family member (HSP81-2) as well as the ROS scavenging gene peroxiredoxin-5 (PRDX5).

The stress response gene HSP81-2 mentioned above is a molecular chaperone involved in cell cycle control and signal transduction. A total of 11 other genes implicated in replication machinery and cell division were also differentially expressed during the infection process (Fig. 4e, 6 and Supplementary Tables S9). Unlike the other functional categories discussed above, in the case of the other 11 genes involved in replication/cell division, the general pattern was of upregulation relative to controls. We interpret these latter changes as implying enhancement of symbiont cell division post-infection, in order to establish an *in hospite* population. Note that Fig 1a is consistent with this hypothesis.

#### Genes implicated in metabolism

Although the exact composition of the translocated material and mechanisms underlying exchange between the coral host and algal symbionts are still unclear (and may vary considerably between associations), a considerable body of evidence points to these being sugars, sterol/lipids and nitrogen compounds (reviewed in Davy et al., 2012), hence one specific focus in the present case is on genes potentially involved in carbohydrate and nitrogen metabolism, and in metabolite transport.

Carbonic anhydrases (CAs) are an essential component of carbon-concentrating mechanisms (CCMs); two genes encoding CAs and, consistent with the hypothesis that sugars are key metabolites in the association, 11 genes involved in carbohydrate metabolism were differentially expressed during the infection process (Fig. 5a, 6 and Table 1). Interestingly, both of the CAs were upregulated during infection, mtCA2 transiently (up in all three replicates at 48h post-infection) and cynT earlier and more consistently. Several genes in this category showed dynamic expression patterns through the infection process, including two genes involved in breaking down stored carbohydrates - the starch debranching enzyme, limit dextrinase (LDA) and the glycogen debranching enzyme AGL. Several genes were strongly downregulated through the infection process relative to controls, including those encoding glyceraldehyde-3-phosphate dehydrogenase (G3P) and fructose bisphosphate aldolase (FBA1) (Fig. 5a, 6 and Table 1). Nine genes implicated in photosynthesis were differentially expressed in *Cladocopium* during the infection process. While the majority of these were downregulated relative to the controls, two genes (psbV and phot2) were upregulated in the later stages of infection. Of these phot2, which encodes the photoreceptor phototropin-2 (phot2) (Supplementary Table S10), is of particular interest given the roles of phototropins in chloroplast movement to maximize light capture under low light conditions (Kagawa et al., 2001).

**Fig. 5.**
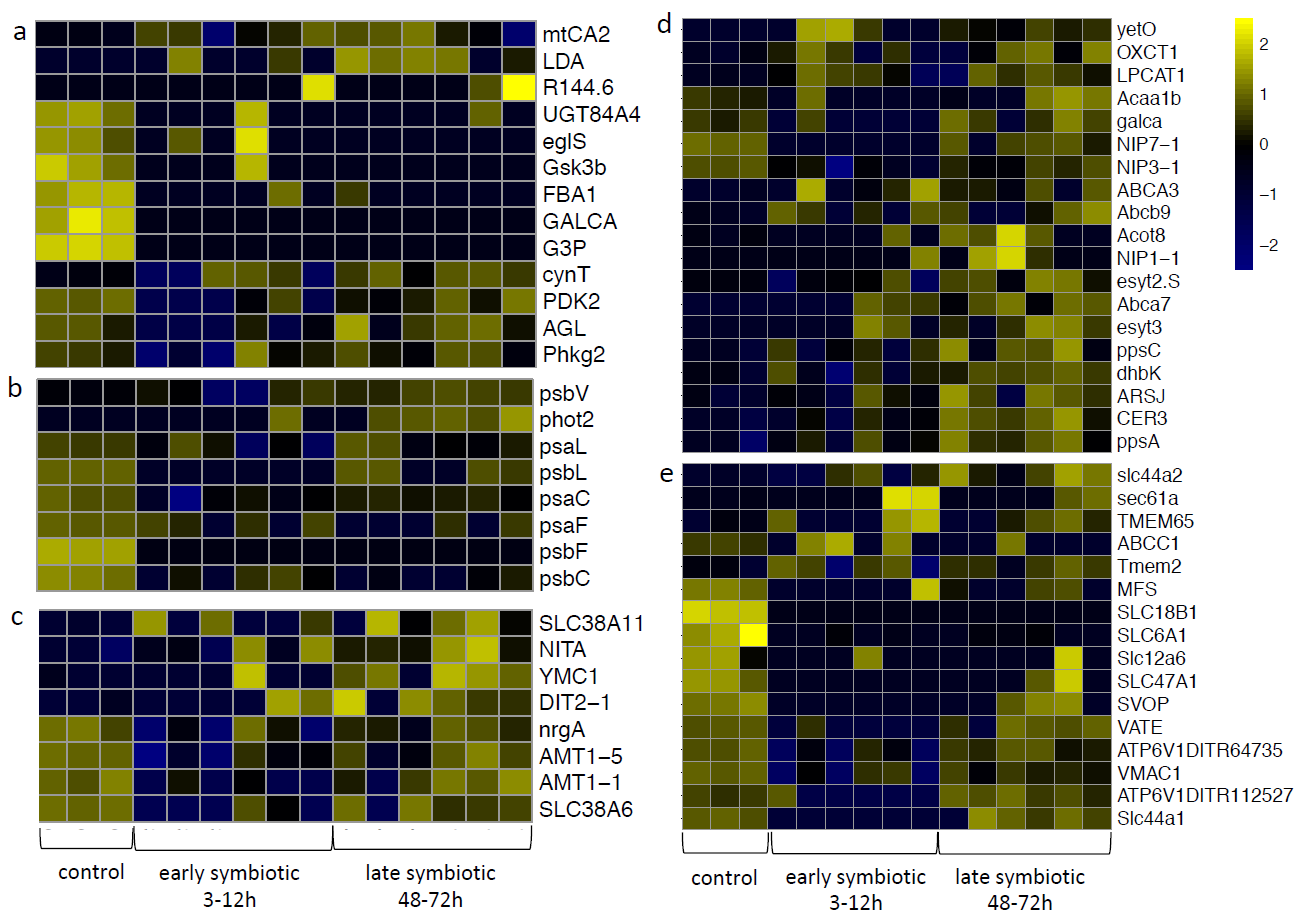
Heat maps of **a** genes implicated in carbohydrate metabolism, **b** genes implicated in photosynthesis, **c** genes implicated in nitrogen metabolism, **d** genes implicated in lipid metabolism, **e** genes encoding transporters in symbiotic *Cladocopium* at 3h, 12h, 48h, 72h post-infection and in controls. The hierarchical clustering shown here was obtained by comparing the expression values (fragments per kilobase of transcript per million; FPKM) for symbiotic samples and the controls. Expression values were log_2_-transformed and median centered by transcript. Relative expression levels are shown in yellow (upregulated) and purple (downregulated).

**Fig. 6.**
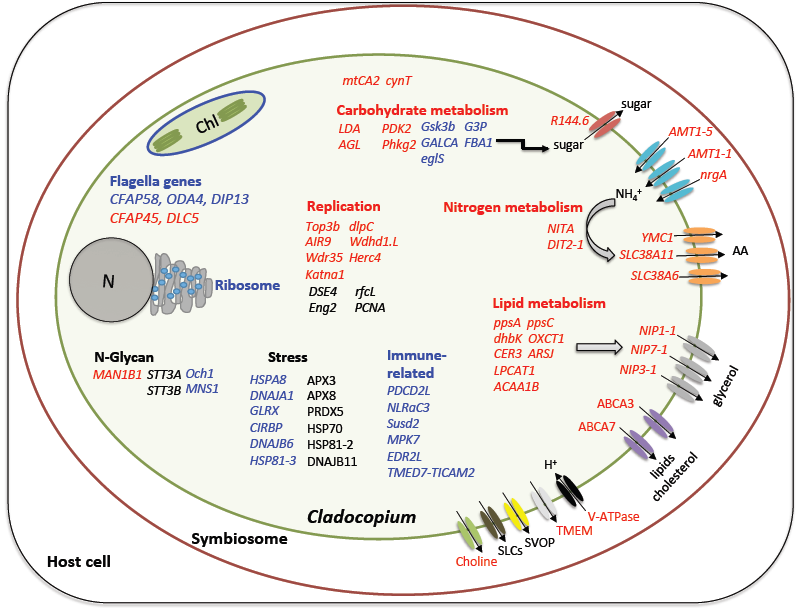
Integrative model of the genes and pathways occurring in *Cladocopium* during lifestyle transition from free-living to symbiotic. The diagram shows algal genes and their roles during the infection of *Acropora tenuis* larvae. Genes in red are upregulated, blue are downregulated, black are differentially regulated throughout the course of the infection. The interaction involved changes in transcriptional level of many genes potentially implicated in changes in lifestyles, including environmental sensing genes (N-Glycan, ANK, AR), flagellar genes, and stress- and immune-related genes. The interaction hallmark is changes in genes involved in carbohydrate, nitrogen and lipid metabolism and transport providing the first evidence of active metabolic efflux from the algal symbiont to the coral host at this early stage of the symbiosis establishment. Expression data for those genes are presented in Fig 4, 5 and Supplementary Table S12.

Eight genes involved in nitrogen metabolism were differentially expressed (Fig. 5c, 6 and Table 1). Four genes that were initially expressed were downregulated until late in the infection process: the ammonium transporters nrgA, AMT1-1 and AMT1-5, and the solute carrier SLC38A6. Conversely four genes that were not significantly expressed in the free-living state were upregulated through the infection process: nitrate reductase (NITA), the glutamate/malate translocator DIT2-1, the solute carrier SLC38A11 and the organic transporter YMC1.

A large number of genes (19) likely to be involved in lipid metabolism and transport were differentially expressed during the course of the infection, seventeen being upregulated in infection and the other two (NIP7-1 and NIP3-1, both encoding aquaporins) being initially downregulated but returning to the free-living levels late in the infection process (Fig. 5d, 6 and Table 1). Among the upregulated DEGs, ATP binding cassette A member 3 (ABCA3) and ATP-binding cassette sub-family A member 7 (ABCA7) are of particular interest in terms of possible involvement in phospholipid and/or cholesterol efflux from cells.

## Discussion

The majority of corals acquire symbionts from the environment through horizontal transmission, whereby the coral hosts in their early life history stages take up symbionts from the environment (Adams et al., 2009; Cumbo et al., 2012). While larvae are not the only coral life history stage at which symbiont acquisition can occur, larval stages provide a unique opportunity to study the initiation of the mutualism. Previous work on this topic has focused almost exclusively on the host side of the relationship, largely due to inadequate genomic and transcriptomic data for Symbiodinaceae. This imbalance is partly addressed here, where the application of high-throughput sequencing methods permitted the detection of thousands of genes that were differentially expressed in the algal symbiont (*Cladocopium*) during the infection of *Acropora tenuis* larvae. In addition to other novel symbiosis-related signals, the symbiont transcriptomic response provides evidence for active metabolite transport from the symbiont.

### Transcriptome repression during the transition from the free-living state to symbiosis

The most obvious implication of the transcriptomic analyses is that the transition to the symbiotic state in *Cladocopium* is marked by suppression of expression of many genes, including those involved in core functions such as translation. This phenomenon is the molecular corollary of adapting to relatively stable intracellular conditions rather than the variable and unpredictable external environment.

In symbiosis, not only are (for example) environmental sensing and stress response requirements reduced, but also functions related to motility are no longer required. Genome reduction is frequently observed in prokaryotes that are in obligate associations and in parasitic eukaryotes (Wolf and Koonin 2013, Brown & Wernegreen 2016, Alonso et al., 2019). Whether genome reduction has occurred in Symbiodiniaceae is a contentious issue; whilst the genomes of those that have been sequenced to date are large (1-5 Gb; Aranda et al. 2016), they are actually much smaller than those of many other dinoflagellates. For example, the genome of *Prorocentrum micans* is ∼250Gb. The fact that extensive genome reduction is not seen in Symbiodiniaceae presumably reflects the fact that most or all members of this group have a free-living stage, requiring the retention of the ability to respond to changing conditions. The dramatic decrease in transcriptome complexity observed here in adjusting to the symbiotic state may be analogous to - or even a precursor of- the genome reduction observed in many obligate associations. Expression of only a limited subset of genes has also been observed in *Durusdinium trenchii* (Bellantuono et al., 2019) when *in hospite* in adult sea anemones (*Exaiptasia pallida),* implying that reduction of transcriptome complexity during the establishment of symbiosis may be a general theme in anthozoan cnidarians.

#### Molecular signature of lifestyle transition in symbiotic Symbiodiniaceae

Comparing symbiotic to free-living Symbiodiniaceae during the infection of coral larvae provides a unique opportunity for understanding necessary transcriptomic adjustments to the symbiotic lifestyle. Mohamed et al. (2016) reported that the coral host at the larval stage (*Acropora digitifera* in that case) undergoes transient transcriptome remodelling during symbiont uptake and concluded that the symbiont infection process requires an active response from the host to recognise/accept the introduced symbiont and not a passive process as was previously postulated (Voolstra et al., 2009). Consequently, Symbiodiniaceae should also undergo specific alterations at the molecular level to adjust to changes in lifestyles. For example, genes encoding ANK, AR and TPR proteins were differentially regulated in symbiotic Symbiodiniaceae. ANK repeats and AR proteins are of particular interest, because proteins of this type are important in enabling bacterial symbionts to evade phagocytosis in their sponge hosts; symbiont-encoded AR proteins are thus key players in enabling the endosymbiosis between bacteria and sponges (Thomas et al., 2010; Nguyen et al., 2014). ANK repeats and AR proteins are much less numerous in bacteria than in eukaryotes, but the genomes of symbiotic bacteria are typically enriched in ANK and AR proteins to the extent that this feature is considered predicative of a symbiotic lifestyle (Jernigan & Bordenstein 2014). It has recently been reported (Liu et al., 2018) that ankyrin repeats are among the most abundant protein domains encoded by four Symbiodiniaceae genomes (*Symbiodinium, Cladocopium, Fugacium* and *Breviolum*), highlighting their potential role in host-symbiont interaction through domain interactions.

Differential expression of genes coding for N-glycan biosynthesis is likely to be significant in the context of the symbiotic lifestyle. A significant number of genes required for N-glycan early processing were downregulated in early infection, whilst a gene encoding mannosidase alpha class 1B member 1 (MAN1B1) was upregulated. These results are consistent with the involvement of algal N-glycans in both recognition and engulfment stages during establishment of symbiosis (Weis et al 2008; Davy et al 2012). In the case of *Exaiptasia* interacting with Symbiodiniaceae, glycans appear to play critical roles in post-recognition “persistence mechanisms” (Parkinson et al., 2018). Symbiodiniaceae algae that are capable of establishing symbiosis have complex repertoires of glycan processing enzymes that differ from their free-living counterparts (Lin et al 2015 and Mohamed et al. 2019, in prep), suggesting that variation in glycoprotein structure may be involved in host recognition specificity.

#### Suppression of flagellar motility and stress-related functions in symbiotic Symbiodiniaceae

In a sense, the interaction studied here, between *Cladocopium* and coral larvae can be viewed as the opposite of what happens during coral bleaching - where symbionts depart their host due to physiological stress. A corollary of this idea is that genes associated with the initial infection process might be informative in assessment of the “health” of colonies of symbiotic corals. On the host side of this interaction, the small GTPases Rab4/5 and Rab7, together with their nucleotide exchange factors, may be considered indicators of the likely stability of the coral / Symbiodiniaceae interaction, because of their likely roles in regulating phagosome maturation (Mohammed et al., 2016). On the Symbiodiniaceae side of the interaction, genes implicated in flagellar motility may be similarly informative.

The differential expression analyses described here indicate that the flagellar motility genes ODA4 and CFAP58 were downregulated throughout the infection process. CFAP58 codes for a cilia- and flagella-associated protein, whereas ODA4 encodes the flagellar outer dynein arm heavy chain beta, which is a force generating protein of eukaryotic cilia and flagella. Suppression of flagellar genes during the establishment of symbiosis is significant, as within their host, Symbiodiniaceae are in the coccoid immobile stage that lacks flagella, whereas the free-living gymnodinioid stage is motile and possesses two flagella (Titlyanov and Titlyanova 2002). In this context, Edge (2007) showed that upregulation of a calmodulin gene associated with flagellar motility occurred in Symbiodiniaceae *in hospite* during thermal stress and suggested that this response might be indicative of the likelihood of symbiosis breakdown as the symbiont prepares to leave an inhospitable host.

#### Metabolic interactions during the establishment of symbiosis in coral larvae

Glucose is thought to be the major photosynthetic product translocated from the algal symbiont to its cnidarian host (Burriesci et al., 2012; Hillyer et al., 2017). Results presented here suggest that, whereas in free-living Symbiodiniaceae fixed carbon in excess of immediate requirements likely accumulates in glycogen and/or starch, in symbiosis the balance shifts to glycogen / starch breakdown; limit dextrinase is upregulated to enable glucose mobilization, glycogen synthase is downregulated and the carbohydrate transporter R144.6 is activated. These results provide the first evidence of active metabolic response from the symbiont to translocate glucose to the coral host at this early stage of symbiosis. Consistent with these results, transcriptomic analyses imply that activation of glycogen biosynthesis is upregulated in cnidarian hosts during colonisation by compatible Symbiodiniaceae (Matthews et al., 2017, Lin et al., 2019,). Moreover, Kopp et al. (2015) demonstrated the translocation of fixed carbon from Symbiodiniaceae to the coral host and visualised glycogen granules in coral tissues.

In addition to carbohydrate transfer, the transcriptomic data presented above are consistent with translocation of lipids and/or sterols from the symbiont to the (cnidarian) host, as implied by previous studies (Davy et al., 2012, Kopp et al., 2015, Bertucci et al., 2015, Matthews et al., 2017, Radecker et al., 2018, Lin et al., 2019). ABC family A transporters have been implicated in the efflux of cellular cholesterol and phospholipids from mammalian cells (Zhao et al., 2012), therefore the upregulation of genes encoding the ATP binding cassette A members 3 (ABCA3) and 7 (ABCA7) observed in symbiosis is consistent with apolipoprotein-mediated phospholipid and/or cholesterol efflux from symbiont cells. Consistent with these results, Bellantuono et al. (2019) reported upregulation of five ABC transporters in *Durusdinium trenchii* (formerly *Symbiodinium* clade D) in symbiosis compared to a free-living control. An ABC transporter is associated with the symbiosome membrane (Peng et al., 2010) and proteomic analyses indicate that several members of this family are associated with the symbiotic condition in *Exaiptasia* (Matthews et al.,2017). ABC transporters may therefore be key facilitators of metabolite exchange on both sides of the cnidarian - Symbiodiniaceae interaction.

With respect to nitrogen metabolism, the implications of the *Cladocopium* gene expression data – and indeed the broader literature - are less clear. During infection, some ammonium transporters are temporarily downregulated, but return to the free-living levels at the end of the infection, and other transport proteins that are initially off are upregulated during the infection, as is nitrate reductase (Fig 5c and Table 1). Both the coral host and Symbiodiniaceae therefore appear to have the ability to incorporate ammonium (Davy et al., 2012), and although the algae account for most of the uptake of ammonium (NH_4_^+^) when this is highly enriched in the environment (Pernice et al., 2012), they are unlikely to be able to do this under “normal” conditions. In response to symbiont infection, cnidarian hosts typically upregulate the high-affinity ammonium assimilation system (glutamine synthase), ensuring that cytoplasmic ammonium levels are very low (e.g. Lin et al., 2019), possibly enabling the host to restrict growth of the symbiont (Davy et al., 2012, Matthews et al., 2017). The upregulation of nitrate reductase during infection, and the nitrate assimilation capacity of symbionts *in hospite* (Pernice et al., 2012), are hard to rationalise, as nitrate is not a normal animal metabolite. One hypothesis to account for this is that nitrate enters the coral cytoplasm under a non-specific mechanism or as a mimic of a “normal” metabolite. Another scenario is that nitrate and/or nitrite may be produced by prokaryotes or other single celled microbes that are intimately associated with the hosts; metagenomic analysis of the *Porites lutea* association gives this idea some credibility (Robbins et al., in press). It is likely that symbiont / host exchanges involving organic nitrogen compounds such as amino acids also occur (Shinzato et al., 2014; Ying et al., 2018), and the upregulation of several solute carrier proteins in symbiotic *Cladocopium* reported here provides the first direct evidence supporting this.

#### Symbiont acquisition at the larval stage - cost or benefit for the host?

Few previous studies have addressed host-symbiont interactions in coral larvae. Based on the transcriptomic work described above, we conclude that in this case symbionts acquired during the larval stage could potentially contribute to host nutrition prior to settlement, however, whether they actually do so is as yet unclear. Kopp et al. (2016) reported that although metabolite translocation occurred from symbiont to host in the case of *Pocillopora damicornis* larvae, such transfer was insignificant in terms of larval nutrition. More recently, Hartmann et al. (2019) showed that infection of *Orbicella faveloata* larvae by the native photosynthetic symbiont is not advantageous, as infected larvae exhibited reduced motility and decreased survival. In mass spawning corals, the larval endoderm is rich in parentally provisioned storage lipids, which are the primary energy source of larvae (Harii et al., 2007). However, this does not rule out the possibility of photosymbionts contributing more significantly to larval nutrition as storage lipids are depleted.

*Acropora* larvae generally acquire their photosymbionts just before or after settlement and metamorphosis rather than during larval stages, but this might simply be a consequence of the higher abundance of Symbiodiniaceae on substrates than in seawater (Cumbo et al., 2012, Littman et al., 2008). The significance of fortuitous acquisition of compatible symbionts during the larval phase might therefore be that it provides newly settled larvae with the advantage of a “head start” in post-settlement life – rather than there being a time lag before symbionts reach a nutritionally-useful density, infected larvae might essentially “hit the ground running”. However, given the enormous genetic diversity of Symbiodiniaceae (LaJeunesse et al., 2018), and the metabolic (Shinzato et al., 2011, Ying et al., 2018) and reproductive (Harrison and Wallace 1990, Okubo et al., 2013) diversity of corals and their relatives, it may be premature to generalise about the significance of larval uptake of symbionts at this time. Nevertheless, the results presented here are the first to document large-scale gene expression profiles of Symbiodiniaceae in a coral host, the transcriptional responses providing new insights into the potential metabolic contributions of Symbiodiniaceae to the coral association.

## Acknowledgements

The research was supported by the Australian Research Council through Grant CE140100020 to DM via the ARC Centre of Excellence for Coral Reef Studies at James Cook University. AM was supported by PhD scholarships provided by the Egyptian Ministry of Higher Education, James Cook University Postgraduate Research Scholarship (JCUPRS) and AIMS@JCU schemes.

The authors gratefully thank the staff of the SeaSim at AIMS for their assistance during the experiment. The authors also would like to thank Victor Beltran at AIMS for supplying the *Cladocopium* culture used in the infection experiment.

## Author contribution

AM and DM conceived and designed the work. AM, AN, and DB carried out the experiment at AIMS. AM, NA, and AM generated the RNA-Seq data. AM analysed the transcriptomic data. AM, CX, and DM interpreted the results. AM, EB and DM wrote the manuscript. All authors read the article and approved the final version.

## Supplementary results/discussion

### Infection dynamics

*Cladocopium* densities increased from 10 cells (mean abundance, n=30) at the first sampling during infecting *Acropora tenuis* larvae (3h post-infection) and reached the maximum of 120 cells (mean abundance, n=30) at the 27h time point (but with higher variability) suggesting that symbiosis has been developing during the time-course experiment (Fig.1). Indeed, such variability has been reported in previous infection experiments (Harii et al., 2009 and Bay et al., 2012). The number of acquired symbiont cells is highly variable among coral species and might differ according to environmental conditions. Cumbo et al. (2012) reported as few as 10 symbiont cells acquired by larvae in acroporids, whereas Harii et al. (2009) found an average of 50-60 symbiont cells. More recently, Mies et al. (2017b) reported 200 acquired symbionts per larva of the coral *Montipora*.

### Symbiont and coral mapping: dual RNA-Seq

Symbiont and coral signals were separated by mapping the raw reads onto their corresponding reference transcriptomes. The read counts that mapped to the *Cladocopium* transcriptome ranged from 176,232 to 1,786,650 (median of 510,210) (Supplementary Figure S1, Supplementary Table S1). Principal component analysis conducted on the normalised *Cladocopium* counts clearly separates the control from the symbiotic *Cladocopium* along PC1 (Supplementary Fig. S2).

### Gene Ontology (GO) analysis reveals suppression of protein synthesis and processing

DEGs were annotated using the GO database and over-representation of GO terms was used to infer functions at the three categories: molecular function (GO-MF), cellular component (GO-CC) and biological process (GO-BP) that were most affected during infection. Using a corrected *P*-value ≤ 0.05, no significant GO enrichment of any of the three categories could be detected among upregulated genes. For downregulated genes, significant enrichment was detected at each time point (Supplementary Table S3). At 3 h downregulated genes showed significant GO enrichment with respect to 11 GO-CC related to ribosome, Golgi, chloroplast and membrane functions and one GO-MF “*structural constituent of ribosome”.* Of these down-regulated genes at 3 h*, “cytosolic large ribosomal subunit”* was the most over-represented GO-CC term. At 12 h downregulated genes showed significant GO enrichment with respect to 3 GO-BP all related to translation, 14 GO-CC related to ribosome, Golgi, chloroplast and membrane functions and one GO-MF “*structural constituent of ribosome”*. Of these downregulated genes at 12 h, “*cytosolic ribosome”* was the most over-represented GO-CC term. At 48 h down-regulated genes showed significant GO enrichment with respect to 10 GO-CC related to ribosome, Golgi, chloroplast and membrane functions and two GO-MF “*structural constituent of ribosome”* and “*mRNA binding”*. At 72 h downregulated genes showed significant GO enrichment with respect to one GO-BP “*translation”*, 13 GO-CC related to ribosome, chloroplast and membrane functions and two GO-MF “*structural constituent of ribosome”* and “mRNA binding”

### Transporter activity

Free-living Symbiodiniaceae are expected to encounter nutrients and ions different from those experienced by symbiotic counterparts. This should be reflected in transporter activity. In symbiotic *Cladocopium,* 16 genes encoding several transporter proteins were differentially expressed in this study (Fig. 5e, 6 and Supplementary Tables S11) including transmembrane proteins, solute carriers (SLCs), and ion transporters. Six genes encoding transmembrane transporters were upregulated including 2 genes encoding transmembrane proteins 65/2 and two genes encoding SV2 related protein (SVOP), which has transmembrane transporter activity. Five genes encoding proteins involved in ion transport were differentially expressed, all of which belong to the vacuolar-type H^+^-ATPase (V-ATPase) protein family. V-ATPase acidifies intracellular organelles by pumping protons across plasma membrane. Six SLC genes showed differential expression, among them 2 genes encoding solute carrier family 44 protein member 2 (SLC44A2) were upregulated. SLC44A2 is a multi-pass membrane protein that might act as a choline transporter.

## Supplementary Tables

**Table S1.**
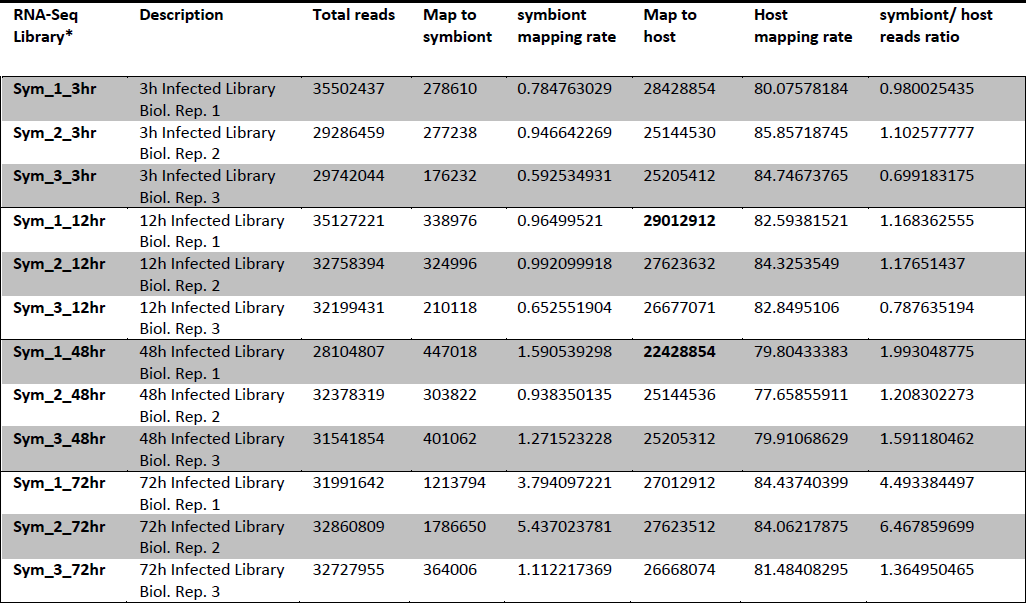
High-throughput sequencing and mapping summary. 12 RNA-Seq libraries were sequenced on an Illumina Hi-Seq platform and produced a total of about 400 million reads from *Cladocopium*-infected *Acropora tenuis* larvae during the time-course infection experiment over 4 time points. There were 3 biological replicates per time point. Reads were separated in silico by mapping to the corresponding transcriptome. The symbiont/host read ratio increased during infection and reached the maximum of 6.5 at 72 h post infection.

Table S2 Identified differentially expressed genes (DEGs) (P ≤ 0.01) in *Cladocopium* during infecting *Acropora tenuis* larvae *Provided as excel file*

**Table S3.**
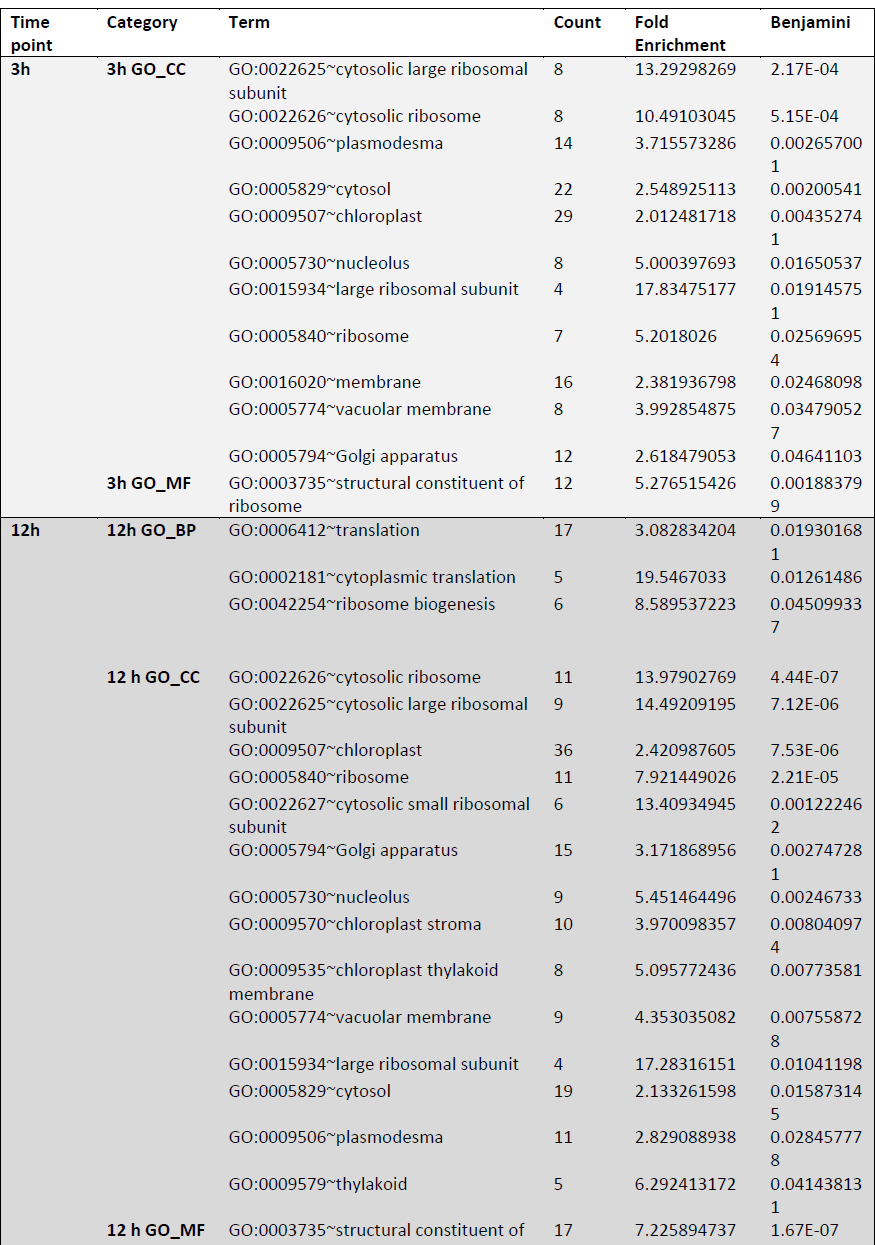
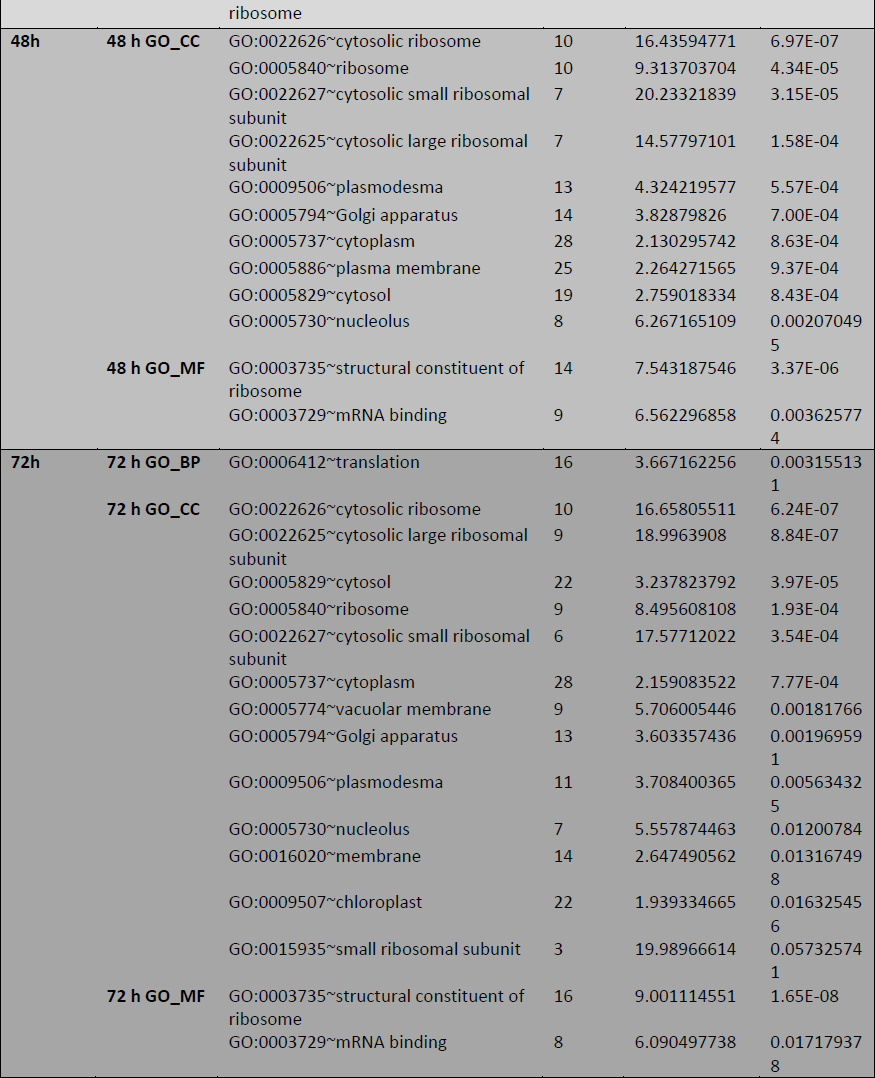
Gene Ontology (GO) categories (GO-BP, CC, MF) enriched (with corrected P ≤ 0.05 among downregulated genes

**Table S4.** Swiss-Prot (SP) annotation of differentially expressed transcripts (P ≤ 0.01) likely involved in N-Glycan biosynthesis in symbiotic *Cladocopium*. E-value threshold of 10^-5^ was applied in BLASTX searches against the SP database

| Transcript ID | Swiss-Prot ID | Annotation |
| --- | --- | --- |
| TR62692\|c0_g1_i3 | Q9UKM7 | mannosidase alpha class 1B member 1(MAN1B1) |
| TR61938\|c0_g2_i1 | Q9UTR6 | alpha-1,6-mannosyltransferase Och1(och1) |
| TR47738\|c0_g1_i2 | Q9C512 | alpha-mannosidase 1(MNS1) |
| TR58977\|c0_g1_i4 | Q9Y719 | alpha-1,3-glucan synthase Mok13(mok13) |
| TR56360\|c0_g1_i1 | Q5RCE2 | STT3A, catalytic subunit of the oligosaccharyltransferase complex (STT3A) |
| TR12916\|c0_g1_i1 | Q8TCJ2 | STT3B, catalytic subunit of the oligosaccharyltransferase complex (STT3B) |
| TR3683\|c0_g2_i1 | Q9XGM8 | alpha-1,3-mannosyl-glycoprotein beta-1,2-N-acetylglucosaminyltransferase (CGL1) |

**Table S5.**
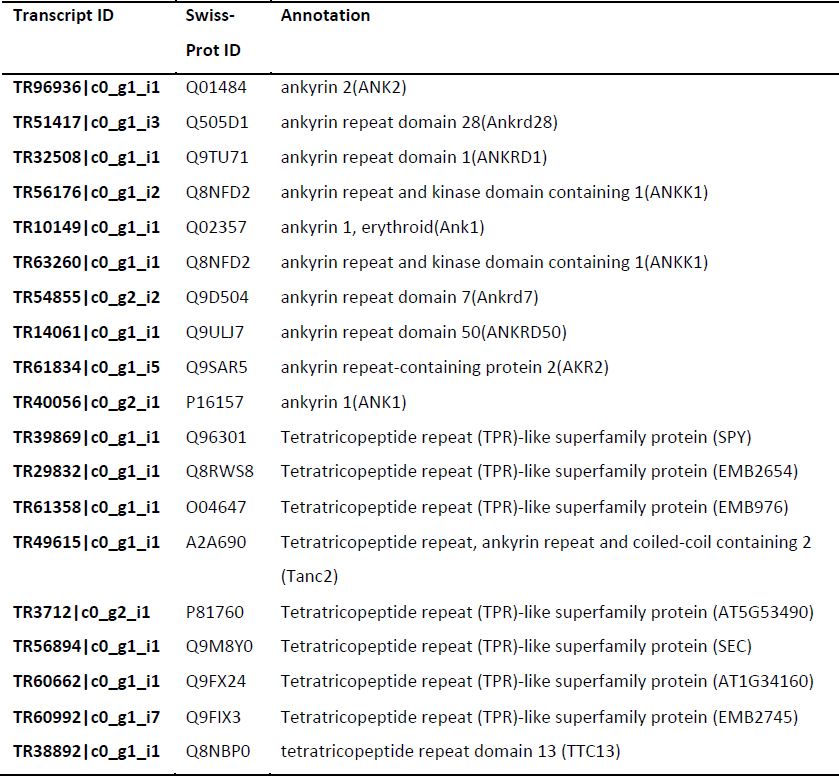
Swiss-Prot (SP) annotation of differentially expressed transcripts (P ≤ 0.01) encoding ankyrin (ANK) proteins, ankyrin repeat (AR) containing proteins and tetratricopeptide repeat (TPR) containing proteins in symbiotic *Cladocopium*. E-value threshold of 10^-5^ was applied in BLASTX searches against the SP database

| Transcript ID | Swiss-Prot ID | Annotation |
| --- | --- | --- |
| TR96936\|c0_g1_i1 | Q01484 | ankyrin 2(ANK2) |
| TR51417\|c0_g1_i3 | Q505D1 | ankyrin repeat domain 28(Ankrd28) |
| TR32508\|c0_g1_i1 | Q9TU71 | ankyrin repeat domain 1(ANKRD1) |
| TR56176\|c0_g1_i2 | Q8NFD2 | ankyrin repeat and kinase domain containing 1(ANKK1) |
| TR10149\|c0_g1_i1 | Q02357 | ankyrin 1, erythroid(Ank1) |
| TR63260\|c0_g1_i1 | Q8NFD2 | ankyrin repeat and kinase domain containing 1(ANKK1) |
| TR54855\|c0_g2_i2 | Q9D504 | ankyrin repeat domain 7(Ankrd7) |
| TR14061\|c0_g1_i1 | Q9ULJ7 | ankyrin repeat domain 50(ANKRD50) |
| TR61834\|c0_g1_i5 | Q9SAR5 | ankyrin repeat-containing protein 2(AKR2) |
| TR40056\|c0_g2_i1 | P16157 | ankyrin 1(ANK1) |
| TR39869\|c0_g1_i1 | Q96301 | Tetratricopeptide repeat (TPR)-like superfamily protein (SPY) |
| TR29832\|c0_g1_i1 | Q8RWS8 | Tetratricopeptide repeat (TPR)-like superfamily protein (EMB2654) |
| TR61358\|c0_g1_i1 | O04647 | Tetratricopeptide repeat (TPR)-like superfamily protein (EMB976) |
| TR49615\|c0_g1_i1 | A2A690 | Tetratricopeptide repeat, ankyrin repeat and coiled-coil containing 2 (Tanc2) |
| TR3712\|c0_g2_i1 | P81760 | Tetratricopeptide repeat (TPR)-like superfamily protein (AT5G53490) |
| TR56894\|c0_g1_i1 | Q9M8Y0 | Tetratricopeptide repeat (TPR)-like superfamily protein (SEC) |
| TR60662\|c0_g1_i1 | Q9FX24 | Tetratricopeptide repeat (TPR)-like superfamily protein (AT1G34160) |
| TR60992\|c0_g1_i7 | Q9FIX3 | Tetratricopeptide repeat (TPR)-like superfamily protein (EMB2745) |
| TR38892\|c0_g1_i1 | Q8NBP0 | tetratricopeptide repeat domain 13 (TTC13) |

**Table S6.** Swiss-Prot (SP) annotation of differentially expressed transcripts (P ≤ 0.01) involved in flagellar motility in symbiotic *Cladocopium*. E-value threshold of 10^-5^ was applied in BLASTX searches against the SP database

| Transcript ID | Swiss-Prot ID | Annotation |
| --- | --- | --- |
| TR63060\|c0_g1_i1 | Q5T655 | cilia and flagella associated protein 58(CFAP58) |
| TR62489\|c0_g1_i1 | Q39565 | flagellar outer dynein arm heavy chain beta(ODA4) |
| TR97365\|c0_g1_i1 | Q39591 | flagellar outer arm dynein 14 kDa light chain LC5(DLC5) |
| TR43719\|c0_g1_i2 | Q9UL16 | cilia and flagella associated protein 45(CFAP45) |
| TR68721\|c0_g1_i1 | Q9XF62 | deflagellation inducible protein(DIP13) |

**Table S7.** Swiss-Prot (SP) annotation of differentially expressed transcripts (P ≤ 0.01) with immune-related functions in symbiotic *Cladocopium*. E-value threshold of 10^-5^ was applied in BLASTX searches against the SP database

| Transcript ID | Swiss-Prot ID | Annotation |
| --- | --- | --- |
| TR39615\|c0_g1_i1 | Q5ZID2 | programmed cell death 2-like (PDCD2L) |
| TR64180\|c0_g2_i8 | Q7RTR2 | NLR family CARD domain containing 3 (NLRC3) |
| TR33508\|c0_g1_i1 | Q9VWW5 | Mitogen-activated protein kinase phosphatase 3 (Mkp3) |
| TR60853\|c0_g1_i2 | Q39027 | MAP kinase 7 (MPK7) |
| TR6597\|c0_g1_i1 | Q9Y3B3 | TMED7-TICAM2 readthrough (TMED7-TICAM2) |
| TR64256\|c0_g3_i1 | Q9DBX3 | sushi domain containing 2 (Susd2) |
| TR43760\|c0_g1_i2 | Q8VZF6 | Pleckstrin homology (PH) domain-containing protein / lipid-binding START domain-containing protein (AT5G45560) |
| TR52242\|c0_g1_i2 | Q5DU56 | NLR family, CARD domain containing 3 (Nlrc3) |
| TR64179\|c1_g3_i5 | Q4LDE5 | sushi, von Willebrand factor type A, EGF and pentraxin domain containing 1 (SVEP1) |
| TR51704\|c0_g1_i1 | Q5DU56 | NLR family, CARD domain containing 3(Nlrc3) |
| TR26819\|c0_g1_i1 | Q3TB82 | pleckstrin homology domain containing, family F (with FYVE domain) member 1 (Plekhf1) |

**Table S8.**
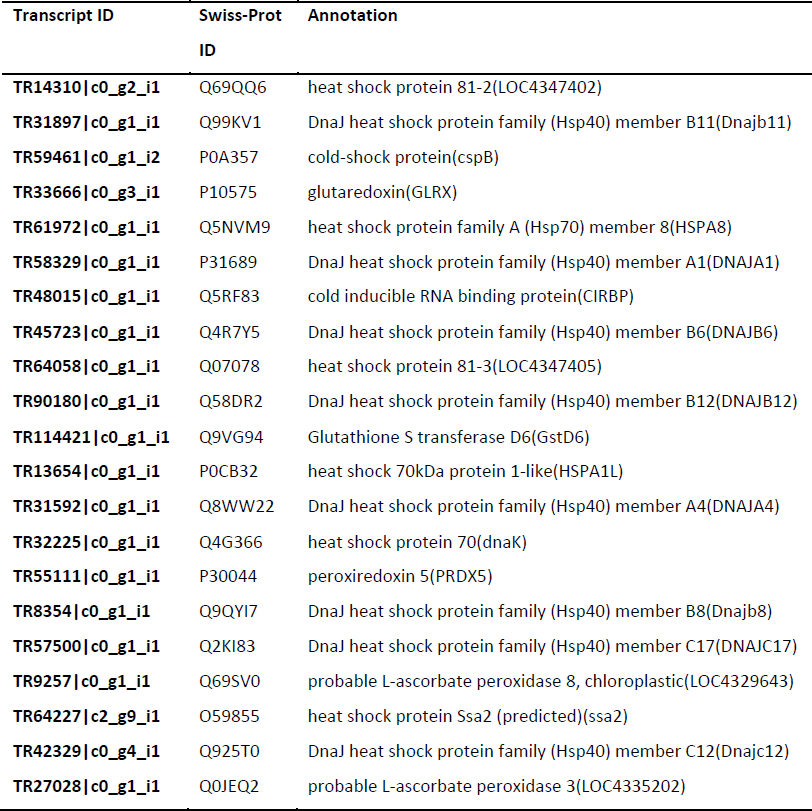
Swiss-Prot (SP) annotation of differentially expressed transcripts (P ≤ 0.01) involved in responses to stress in symbiotic *Cladocopium*. E-value threshold of 10^-5^ was applied in BLASTX searches agains

**Table S9.**
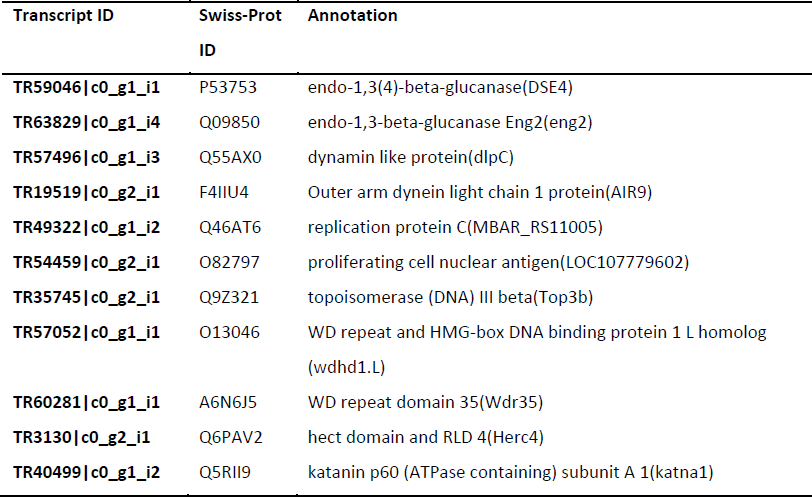
Swiss-Prot (SP) annotation of differentially expressed transcripts (P ≤ 0.01) involved in replication and cell division in symbiotic *Cladocopium*. E-value threshold of 10^-5^ was applied in BLASTX searches against the SP database

**Table S10.** Swiss-Prot (SP) annotation of differentially expressed transcripts (P ≤ 0.01) involved in photosynthesis in symbiotic *Cladocopium*. E-value threshold of 10^-5^ was applied in BLASTX searches against the SP database

| Transcript ID | Uniprot ID | Best BLAST Hit |
| --- | --- | --- |
| TR57075\|c0_g1_i4 | Q85FP8 | photosystem I reaction center subunit XI(psaL) |
| TR61769\|c0_g1_i4 | P51193 | photosystem I subunit III(psaF) |
| TR58592\|c0_g1_i3 | A0T0C6 | photosystem II cytochrome c550(psbV) |
| TR35567\|c0_g1_i2 | Q1XD97 | photosystem II reaction center subunit VI(psbF) |
| TR30415\|c0_g1_i1 | P49472 | PSII 43 kDa protein(psbC) |
| TR57052\|c1_g1_i4 | Q4G382 | PSII reaction center subunit XII(psbL) |
| TR99055\|c0_g1_i1 | A0T0L2 | photosystem i iron-sulfur center |
| TR32083\|c0_g1_i2 | Q9ST27 | phototropin-2(phot2) |

**Table S11.**
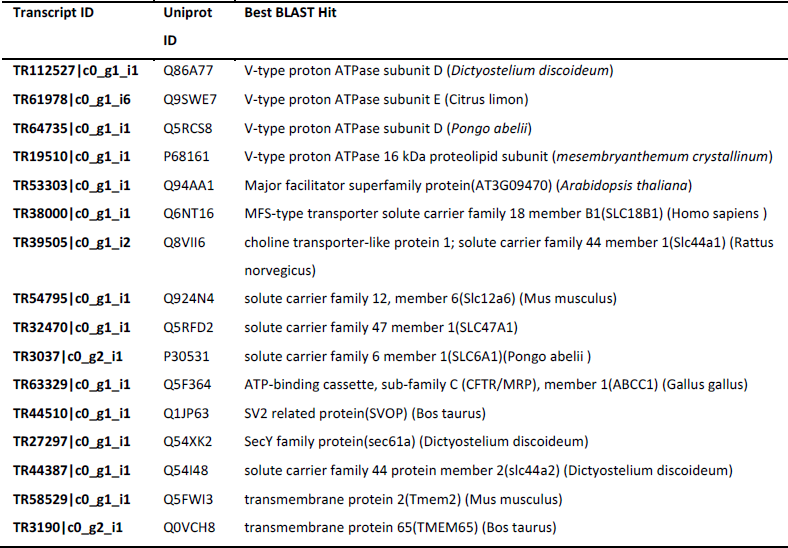
Swiss-Prot (SP) annotation of differentially expressed transcripts (P ≤ 0.01) coding for transporters in symbiotic *Cladocopium*. E-value threshold of 10^-5^ was applied in BLASTX searches against the SP database

Table S12 Normalized expression values for all DEGs discussed in the paper *Provided as excel file*

## Supplementary Figures

**Figure S1.**
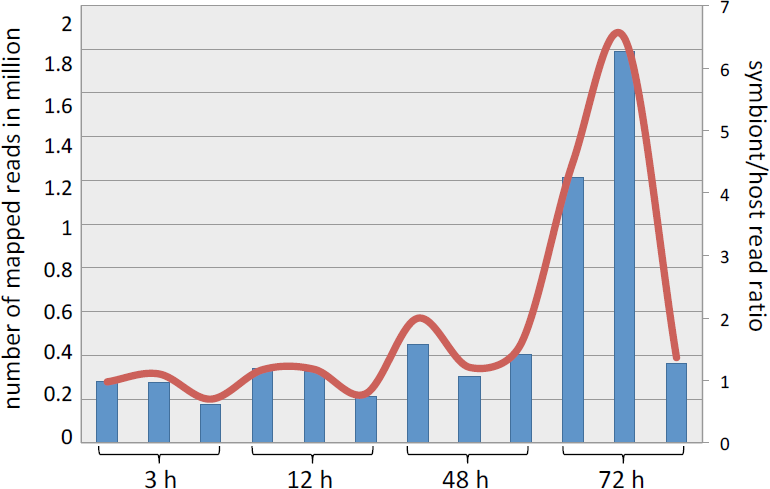
Symbiont and host dual RNA-Seq. Temporal change in transcriptional output was estimated from symbiont/host ratio of total read counts, assuming that the host mRNA output is constant over time. The blue bars represent the number of mapped symbiont reads while the red line represents ratio of symbiont/host mapped reads during the course of infection.

**Figure S2.**
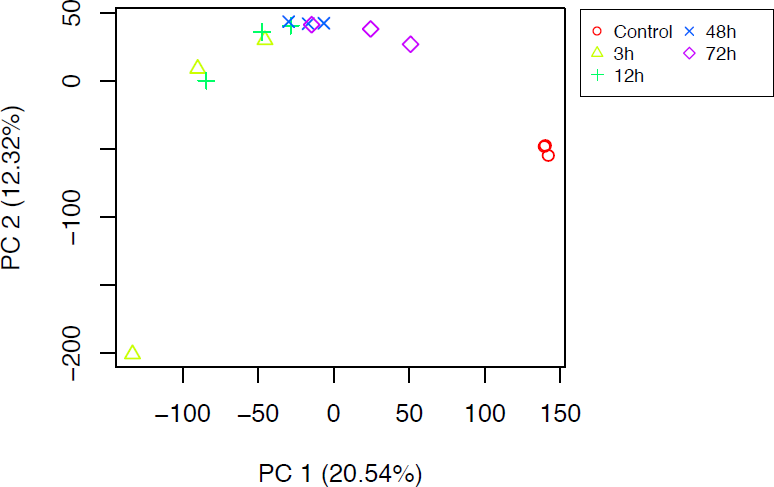
Principal Component Analysis (PCA) plot of log-transformed and normalized count data for triplicates of the four time points post *Cladocopium* infection and the triplicates of the *Cladocopium* control condition.

**Figure S3.**
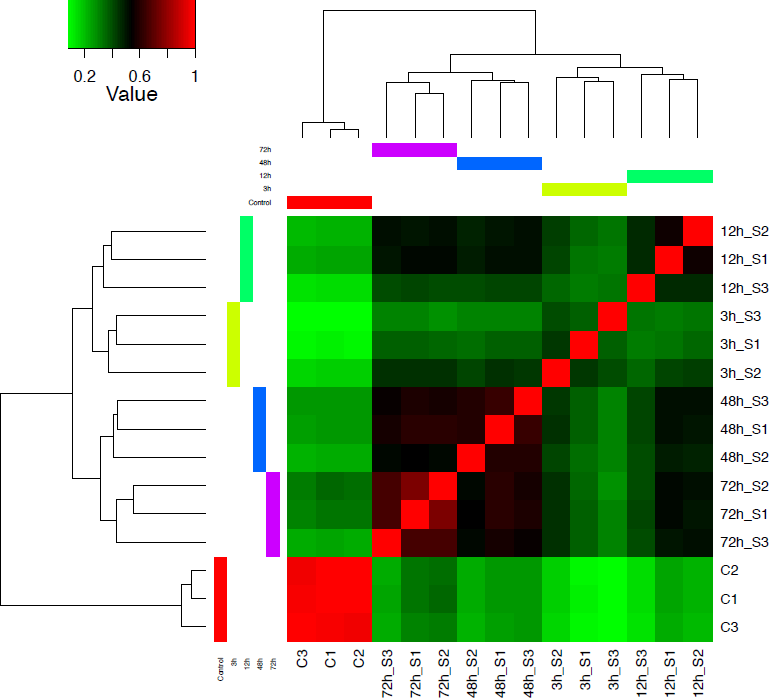
Level of agreement among the biological replicates during *Cladocopium* infection. The heat map shows the hierarchically clustered Spearman correlation matrix resulting from comparing the transcript expression values (TMM-normalized FPKM) for all samples against one another. Sample clustering indicates the consistency between the biological triplicates of the *Cladocopium* infection at each of the 4 time points (3, 12, 48 and 72h) and the triplicate of the *Cladocopium* control condition. The level of correlation is presented by a color field that ranges from green (correlation coefficient 0.2) to red (correlation coefficient 1.0).

## References

Adams LM, Cumbo VR, Takabayashi M. Exposure to sediment enhances primary acquisition of Symbiodinium by asymbiotic coral larvae. Mar Ecol Prog Series. 2009; 377:149–156.

Allemand D, Furla P, Bénazet-Tambutté S. Mechanisms of carbon acquisition for endosymbiont photosynthesis in Anthozoa. Can J of Botany. 1998; 7:925–941.

Alonso DP, Mancini MV, Damiani C, Cappelli A, Ricci I, Alvarez MVN, et al. Genome Reduction in the Mosquito Symbiont *Asaia*. Genome Biol Evol. 2019; 11:1–10.

Aranda M, Li Y, Liew YJ, Baumgarten S, Simakov O, Wilson MC., et al. Genomes of coral dinoflagellate symbionts highlight evolutionary adaptations conducive to a symbiotic lifestyle. Sci Reports. 2016; 6:39734.

Bellantuono AJ, Dougan KE, Granados-Cifuentes C, Rodriguez-Lanetty M. Transcriptome landscape of a thermal-tolerant coral endosymbiont reveals molecular signatures of symbiosis and dysbiosis. BioRxiv. 2019; 508184. https://doi.org/10.1101/508184

Bertucci A, Foret S, Ball EE, Miller DJ. Transcriptomic differences between day and night in Acropora millepora provide new insights into metabolite exchange and light-enhanced calcification in corals. Mol Ecol. 2015; 24:4489–4504.

Bertucci A, Zoccola D, Tambutté S, Vullo D, Supuran CT. Carbonic anhydrase activators. The first activation study of a coral secretory isoform with amino acids and amines. Bioorg Med Chem. 2010;18:2300–2303.

Brown BP, Wernegreen JJ. Deep divergence and rapid evolutionary rates in gut-associated Acetobacteraceae of ants. BMC Microbiol. 2016; 16:140.

Bucher M, Wolfowicz I, Voss PA, Hambleton EA, Guse A. Development and symbiosis establishment in the cnidarian endosymbiosis model Aiptasia sp. Sci Reports. 2016; 6:19867.

Burriesci MS, Raab TK, Pringle JR. Evidence that glucose is the major transferred metabolite in dinoflagellate–cnidarian symbiosis. J Exp Biol. 2012; 215:3467–3477.

Choi YJ, Aliota MT, Mayhew GF, Erickson SM, Christensen BM. Dual RNA-seq of parasite and host reveals gene expression dynamics during filarial worm–mosquito interactions. PLoS Neglected Tropical Diseases. 2014; 8:e2905.

Cumbo VR, Baird AH, Van Oppen MJ. The promiscuous larvae: flexibility in the establishment of symbiosis in corals. Coral Reefs. 2013;32:111–20.

Davy SK, Allemand D, Weis VM. Cell biology of cnidarian-dinoflagellate symbiosis. Microbiol Mol Biol Rev. 2012; 76:229–261.

Edge S. Using Microarrays to Quantify Stress Responses in Natural Populations of Coral. PhD Dissertation. Georgia Institute of Technology. August, 2007.

Grant AJ, Rémond M, People J, Hinde R. Effects of host tissue homogenate of the scleractinian coral Plesiastrea versipora on glycerol metabolism in isolated symbiotic dinoflagellates. Mar Biol. 1997;128:665–670.

Haas BJ, Papanicolaou A, Yassour M, Grabherr M, Blood PD, Bowden J, et al. De novo transcript sequence reconstruction from RNA-seq using the Trinity platform for reference generation and analysis. Nature Protocols. 2013;8:1494–512.

Harii S, Nadaoka K, Yamamoto M, Iwao, K. Temporal changes in settlement, lipid content and lipid composition of larvae of the spawning hermatypic coral Acropora tenuis. Mar Ecol Prog Series. 2007;346:89–96.

Harrison PL, Wallace CC, 1990. Reproduction, dispersal and recruitment of scleractinian corals. In: Dubinsky, Z. (ed.), Ecosystems of the World: Coral Reefs. Elsevier, Amsterdam, Netherlands, 1990, pp 133–207.

Hartmann AC., Marhaver KL, Klueter A, Lovci MT, Closek CJ, Diaz E, et al. Acquisition of obligate mutualist symbionts during the larval stage is not beneficial for a coral host. Mol Ecol. 2019;28:141–155.

Hillyer KE, Dias DA, Lutz A, Roessner U, Davy SK. Mapping carbon fate during bleaching in a model cnidarian symbiosis: the application of 13C metabolomics. New Phytol. 2017;214:1551–1562.

Jernigan KK, Bordenstein SR. Ankyrin domains across the Tree of Life. PeerJ. 2014;2:e264.

Kagawa T, Sakai T, Suetsugu N, Oikawa K, Ishiguro S, Kato T, et al. Arabidopsis NPL1: a phototropin homolog controlling the chloroplast high-light avoidance response. Science. 2001;291:2138–2141.

Kopp C, Domart-Coulon I, Escrig S, Humbel BM, Hignette M, Meibom A. Subcellular investigation of photosynthesis-driven carbon assimilation in the symbiotic reef coral Pocillopora damicornis. MBio. 2015;6:e02299–14.

Kopp C, Domart-Coulon I, Barthelemy D, Meibom A. Nutritional input from dinoflagellate symbionts in reef-building corals is minimal during planula larval life stage. Science Advances. 2016;2:e1500681.

LaJeunesse TC, Parkinson JE, Gabrielson PW, Jeong HJ, Reimer JD, Voolstra CR, et al. Systematic revision of Symbiodiniaceae highlights the antiquity and diversity of coral endosymbionts. Curr Biol. 2018;28:2570–2580

LaMonte G, Orjuela-Sanchez P, Calla J, Wang L, Li S, Swann J, et al. Dual rnaseq shows the human mucosal immunity protein, muc13, is a hallmark of Plasmodium exoerythrocytic infection. Nat Commun. 2019; 10:488

Langmead B, Salzberg SL. Fast gapped-read alignment with Bowtie 2. Nature Methods. 2012;9:357.

Leggat W, Buck BH, Grice A, Yellowlees D. The impact of bleaching on the metabolic contribution of dinoflagellate symbionts to their giant clam host. Plant Cell Environ. 2003; 26:1951–1961.

Levin RA, Beltran VH, Hill R, Kjelleberg S, McDougald D, Steinberg PD, et al. Sex, scavengers, and chaperones: transcriptome secrets of divergent Symbiodinium thermal tolerances. Mol Biol Evol. 2016;33:2201–2215.

Li B, Dewey CN. RSEM: accurate transcript quantification from RNA-Seq data with or without a reference genome. BMC Bioinformatics. 2011;12:323.

Lin K-L, Wang J-T, Fang L-S Participation of glycoproteins in zooxanthella cell walls in the establishment of a symbiotic relationship with the sea anemone, *Aiptasia pulchella*. Zool Stud. 2000; 39:172–178.

Lin S, Cheng S, Song B, Zhong X, Lin X, Li W, et al. The Symbiodinium kawagutii genome illuminates dinoflagellate gene expression and coral symbiosis. Science. 2015;350:691–694.

Lin MF, Takahashi S, Forêt S, Davy SK, Miller DJ. Transcriptomic analyses highlight the likely metabolic consequences of colonization of a cnidarian host by native or non-native Symbiodinium species. Biology Open. 2019;8:bio038281.

Littman RA, van Oppen MJH, Willis BL. Methods for sampling free-living Symbiodinium (zooxanthellae) and their distribution and abundance at Lizard Island (Great Barrier Reef). J Exp Mar Biol Ecol. 2008;364:48–53.

Liu H, Stephens TG, González-Pech RA, Beltran VH, Lapeyre B, Bongaerts P. Symbiodinium genomes reveal adaptive evolution of functions related to coral-dinoflagellate symbiosis. Comm. Biol. 2018;1:95.

Matthews JL, Crowder CM, Oakley CA, Lutz A, Roessner U, Meyer E, et al. Optimal nutrient exchange and immune responses operate in partner specificity in the cnidarian-dinoflagellate symbiosis. Proc Natl Acad Sci USA. 2017;114:13194–13199.

Mies M, Sumida PYG, Rädecker N, Voolstra CR. Marine invertebrate larvae associated with Symbiodinium: a mutualism from the start? Front. Physiol. 2017a; 5:56.

Mies M, Voolstra CR, Castro CB, Pires DO, Calderon EN, Sumida, PYG. Expression of a symbiosis-specific gene in Symbiodinium type A1 associated with coral, nudibranch and giant clam larvae. Roy Soc Open Sci. 2017b;4:170253.

Mohamed AR, Cumbo V, Harii S, Shinzato C, Chan CX, Ragan MA, et al. The transcriptomic response of the coral Acropora digitifera to a competent Symbiodinium strain: the symbiosome as an arrested early phagosome. Mol Ecol. 2016; 25:3127–3141.

Mohamed AR, Cumbo VR, Harii S, Shinzato C, Chan CX, Ragan MA, et al. Deciphering the nature of the coral–Chromera association. ISMEJ. 2018;12:776.

Mohamed AR, Chan CX, Ragan MA, Zhang J, Cooke I, Ball EE, et al. (in preparation) Unravelling the secrets of coral-*Chromera* symbiosis through comparative transcriptomics.

Muscatine L. The role of symbiotic algae in carbon and energy flux in reef corals. In: Dubinsky, Z. (ed.), Ecosystems of the World: Coral Reefs. Elsevier, Amsterdam, Netherlands, 1990, pp. 75–87.

Nguyen MT, Liu M, Thomas T. Ankyrin-repeat proteins from sponge symbionts modulate amoebal phagocytosis. Mol Ecol. 2014;23:1635–1645.

Nuss AM, Beckstette M, Pimenova M, Schmühl C, Opitz W, Pisano F, et al. Tissue dual RNA-seq allows fast discovery of infection-specific functions and riboregulators shaping host–pathogen transcriptomes. Proc Natl Acad Sci USA. 2017;114:E791–800.

Okubo N, Mezaki T, Nozawa Y, Nakano Y, Lien YT, Fukami H, et al. Comparative embryology of eleven species of stony corals (Scleractinia). PLoS One. 2013; 8:e84115.

Papina M, Meziane T, Van Woesik R. Symbiotic zooxanthellae provide the host-coral *Montipora digitata* with polyunsaturated fatty acids. Comp Biochem Physiol B: Biochem Mol Biol. 2003;135:533–537.

Parkinson JE, Tivey TR, Mandelare PE, Adpressa DA, Loesgen S, Weis VM. Subtle differences in symbiont cell surface glycan profiles do not explain species-specific colonization rates in a model cnidarian-algal symbiosis. Front Microbiol. 2018; doi 10.3389/fmicb.2018.00842

Peng S.-E, Wang Y-B, Wang L-H, Chen W-N U, Lu C-Y, Fang L-S, et al. Proteomic analysis of symbiosome membranes in Cnidaria-dinoflagellate endosymbiosis. Proteomics. 2010;10:1002–1016.

Pernice M, Meibom A, Van Den Heuvel A, Kopp C, Domart-Coulon I, Hoegh-Guldberg O, et al. A single-cell view of ammonium assimilation in coral– dinoflagellate symbiosis. ISMEJ. 2012;6:1314.

Rädecker N, Raina JB, Pernice M, Perna, G, Guagliardo P, Kilburn MR, et al. Using Aiptasia as a model to study metabolic interactions in cnidarian-Symbiodinium symbioses. Front Physiol. 2018;9:214.

Robbins S, Singleton C, Chan C, Ying H, Baker A, Messer L, et al. A genomic view of the coral holobiont. In press to: Nature Microbiol.

Robinson MD, McCarthy DJ, Smyth GK. edgeR: a Bioconductor package for differential expression analysis of digital gene expression data. Bioinformatics. 2010;26:139–140.

Robinson MD, Oshlack A. A scaling normalization method for differential expression analysis of RNA-seq data. Genome Biol. 2010;11:R25.

Rosic N, Kaniewska P, Chan CKK, Ling EYS, Edwards D, Dove S, et al. Early transcriptional changes in the reef-building coral Acropora aspera in response to thermal and nutrient stress. BMC Genomics. 2014;15:1052.

Schnitzler CE, Weis VM. Coral larvae exhibit few measurable transcriptional changes during the onset of coral-dinoflagellate endosymbiosis. Mar Genomics. 2010;3:107–116.

Shinzato C, Shoguchi E, Kawashima T, Hamada M, Hisata K, Tanaka M. et al. Using the Acropora digitifera genome to understand coral responses to environmental change. Nature. 2011;476:320.

Shinzato C, Inoue M, Kusakabe M. A snapshot of a coral “holobiont”: a transcriptome assembly of the scleractinian coral, Porites, captures a wide variety of genes from both the host and symbiotic zooxanthellae. PLoS One. 2014; 9:e85182.

Shoguchi E, Beedessee G, Tada I, Hisata K, Kawashima T, Takeuchi T, et al. Two divergent Symbiodinium genomes reveal conservation of a gene cluster for sunscreen biosynthesis and recently lost genes. BMC Genomics. 2018;19:458.

Shoguchi E, Shinzato, C, Kawashima T, Gyoja F, Mungpakdee S, Koyanagi R, et al. Draft assembly of the Symbiodinium minutum nuclear genome reveals dinoflagellate gene structure. Curr Biol. 2013;23:1399–1408.

Thomas T, Rusch D, DeMaere MZ, Yung PY, Lewis M, Halpern A, et al. Functional genomic signatures of sponge bacteria reveal unique and shared features of symbiosis. ISMEJ. 2010;4:1557.

Titlyanov EA, Titlyanova TV. Reef-building corals—symbiotic autotrophic organisms: 2. Pathways and mechanisms of adaptation to light. Russ J Mar Biol. 2002;28:S16–S31.

Trench RK. The cell biology of plant-animal symbiosis. Ann Rev Plant Physiol. 1979;30:485–531.

Voolstra CR, Schwarz JA, Schnetzer J, Sunagawa S, Desalvo MK, Szmant AM, et al. The host transcriptome remains unaltered during the establishment of coral–algal symbioses. Mol Ecol. 2009;18:1823–1833.

Weis VM, Davy SK, Hoegh-Guldberg O, Rodriguez-Lanetty M, Pringle JR. Cell biology in model systems as the key to understanding corals. Trends Ecol Evol. 2008;23:369–376.

Wolf YI, Koonin EV. Genome reduction as the dominant mode of evolution. Bioessays. 2013;35:829–837.

Wolfowicz I, Baumgarten S, Voss PA, Hambleton EA, Voolstra CR, Hatta M, et al. Aiptasia sp. larvae as a model to reveal mechanisms of symbiont selection in cnidarians. Sci Reports. 2016;6:32366.

Wood-Charlson E, Hollingsworth L, Krupp D, Weis V. Lectin/glycan interactions play a role in recognition in coral/dinoflagel-late symbiosis. Cell Microbiol. 2006;8:1985–1993.

Ying H, Cooke I, Sprungala S, Wang W, Hayward DC, Tang Y, et al. Comparative genomics reveals the distinct evolutionary trajectories of the robust and complex coral lineages. Genome Biol. 2018;19:175.

Zhao GJ, Yin K, Fu YC, Tang CK. The interaction of ApoA-I and ABCA1 triggers signal transduction pathways to mediate efflux of cellular lipids. Mol Med 2012;18:149–158.

